# Tracking tRNA evolution in closely related mouse strains

**DOI:** 10.1101/2025.08.30.673268

**Authors:** Andrew D. Holmes, Upasna Sharma

## Abstract

Transfer RNAs are conserved RNAs that are essential for translation in all forms of life and are present in multiple variants to allow interpretation of the entire genetic code. Transfer RNA genes in eukaryotes exist in multiple identical and non-identical copies across the genome to contribute to the tRNA repertoire of a cell. While the same tRNAs are conserved across species, their number and conservation vary from genome to genome. The evolution of these tRNA genes, especially in mammals, is poorly understood. To understand what guides tRNA evolution, we examined the tRNA gene complement from 17 closely related mouse strain genomes. Our analysis revealed that although most tRNAs are conserved, a subset shows copy number variation, sequence divergence, and even strain-specific gains or losses. We also identified a group of transcribed single-copy tRNAs that are conserved across multiple mouse strains and other mammals, suggesting specialized or lineage-specific functions. Together, these findings provide new insights into the evolutionary dynamics of the mammalian tRNA repertoire.

## Introduction

Transfer RNAs (tRNAs) are essential components of the translation machinery, linking codons to amino acids and thereby ensuring accurate and efficient protein synthesis. Beyond their canonical role in translation, tRNAs and their cleavage product, known as tRNA fragments or tRNA-derived RNA (tDRs) ^1^, are increasingly recognized as regulators of gene expression, stress responses, and developmental processes ^2,3^. Given their central role in cellular physiology, tRNA genes are often assumed to be highly conserved across mammalian genomes ^4-6^. However, the true extent of tRNA gene conservation, copy number variation, and transcriptional activity across closely related species or strains has not been systematically explored.

The structure of cytosolic tRNA is conserved in all domains of life. tRNAs generally form a “cloverleaf” secondary structure, with the acceptor arm, the D arm, the anticodon arm, the T arm, and, in some tRNAs, a variable loop ^7,8^. This well-defined, conserved structure of tRNAs allowed numbering all specific bases using the “Sprinzl” numbering system, which can then be used across tRNAs to identify specific bases that are modified, recognized by tRNA-aminoacylases, or differ in sequences ^9^. Moreover, tRNA genes are predicted in assembled genomes based on their sequence and structure being similar to known tRNAs, as established with covariance models ^10^. These tools are used to predict both active tRNA genes and tRNA-like elements. The distinction between the two is primarily based on their sequence and secondary structure similarity to the canonical tRNA models, as assessed by a score generated from a covariance model. This score thus allows for tRNA genes to be ranked by their best-fit and can be used to differentiate inactive tRNA-like elements from transcribed and active tRNAs with some accuracy. These tRNA-like elements can include pseudo-tRNAs, tRNA-like SINE elements, and mitochondrial tRNAs copied into the nuclear genome. Covariance models can also be used to identify tRNAs by amino acids, which, even outside the anticodon sequence, have distinct profiles for each amino acid ^11^.

tRNA genes are transcribed by RNA polymerase III (Pol III) using internal promoters with conserved A box and B box sequences ^12^. However, these sequence features are not sufficient to dictate tRNA expression, as identical tRNA copies can be active or inactive ^13^. While the full determinants of tRNA gene activity are not known, the local genomic context, such as presence in tRNA arrays and CpG density, in combination with internal sequences, provides specificity ^14^. One way to identify transcribed tRNAs is by measuring Pol III binding at tRNA genes ^13^. Moreover, tRNA expression can be regulated by Maf1, a repressor of Pol III ^15^. In addition, tRNAs can be expressed in a tissue-specific manner, although the factors contributing to this specificity remain unclear. For example, tRNA-Arg-TCT-4 is highly expressed in the brain ^16,17^.

Notably, tRNAs exist in many copies in any given eukaryotic genome, allowing for greater transcript redundancy and possibly translation-independent functions. These tRNA sequences show conservation both between orthologs in different species and within, with genomes containing multiple identical copies of certain tRNAs ^18^. While the tRNA genes themselves are highly conserved and often present in multiple copies within genomes, the flanking sequences incorporated into pre-tRNA transcripts show substantial divergence across orthologs and paralogs ^14^. Importantly, tRNAs are also known to vary in copy number among individual humans, despite being highly conserved ^19^. It has been suggested that the maintenance of multiple identical tRNA gene copies across the genome, beyond what may be necessary for function, could result from a process of concerted evolution. According to this idea, which has also been suggested for ribosomal RNA ^20^, the tRNA genes spread across the genome are kept identical in sequence through gene conversion. While this has been induced in yeast tRNAs ^21^, little is known about this process.

Here, we analyze predicted tRNA genes and tRNA-like elements across 17 mouse strains, integrating covariance scoring with Pol III ChIP-seq data to identify actively transcribed tRNAs. We show that while the majority of tRNAs are conserved, subsets of tRNAs exhibit copy number differences, sequence divergence, and even strain-specific gain or loss. Moreover, we identify a set of deeply conserved single-copy tRNAs that are transcribed across multiple strains and mammals, implicating them in specialized and possibly lineage-specific functions. Together, these findings provide new insights into the evolutionary dynamics of the mammalian tRNA repertoire and establish a framework for linking tRNA variation to functional consequences in translation and beyond.

## Results

### RNA polymerase III binding analysis confirms active tRNA gene predictions

To study the number and conservation of tRNA genes across mouse strains, we retrieved the set of predicted tRNAs and tRNA-like elements from the assembled genomes of 17 mouse strains. This set includes 129S1/SvImJ, A/J, AKR/J, CAST/EiJ, CBA/J, DBA/2J, FVB/NJ, NOD/ShiLtJ, NZO/HlLtJ, PWK/PhJ, CBA/J, WSB/EiJ, SPRET/EiJ, C57BL/6NJ, BALB/cJ, IP/J, and C3H/HeJ strain assemblies. Mouse genomes have numerous predicted tRNA-like elements annotated in the genomic tRNA database (gtRNAdb)^22^. These elements, predicted based on their sequence, are either active tRNAs or sequences that partially match active tRNA genes. In addition to tRNAs and pseudo-tRNAs, this set of tRNA-like elements is known to be dominated by tRNA-like SINE elements in the mouse genome ^23^. tRNA-like SINE elements are abundant in rodent genomes compared to the human genome and could cause the count of tRNA-like elements to vary wildly between strains. Our analysis revealed that the total counts of tRNA genes and tRNA-like elements for a given type of tRNA remain broadly the same across the mouse strains, with Ser, Leu, and Asn isotype-derived tRNA elements being most abundant across all strains (**Figure 1A**).

**Figure 1.**
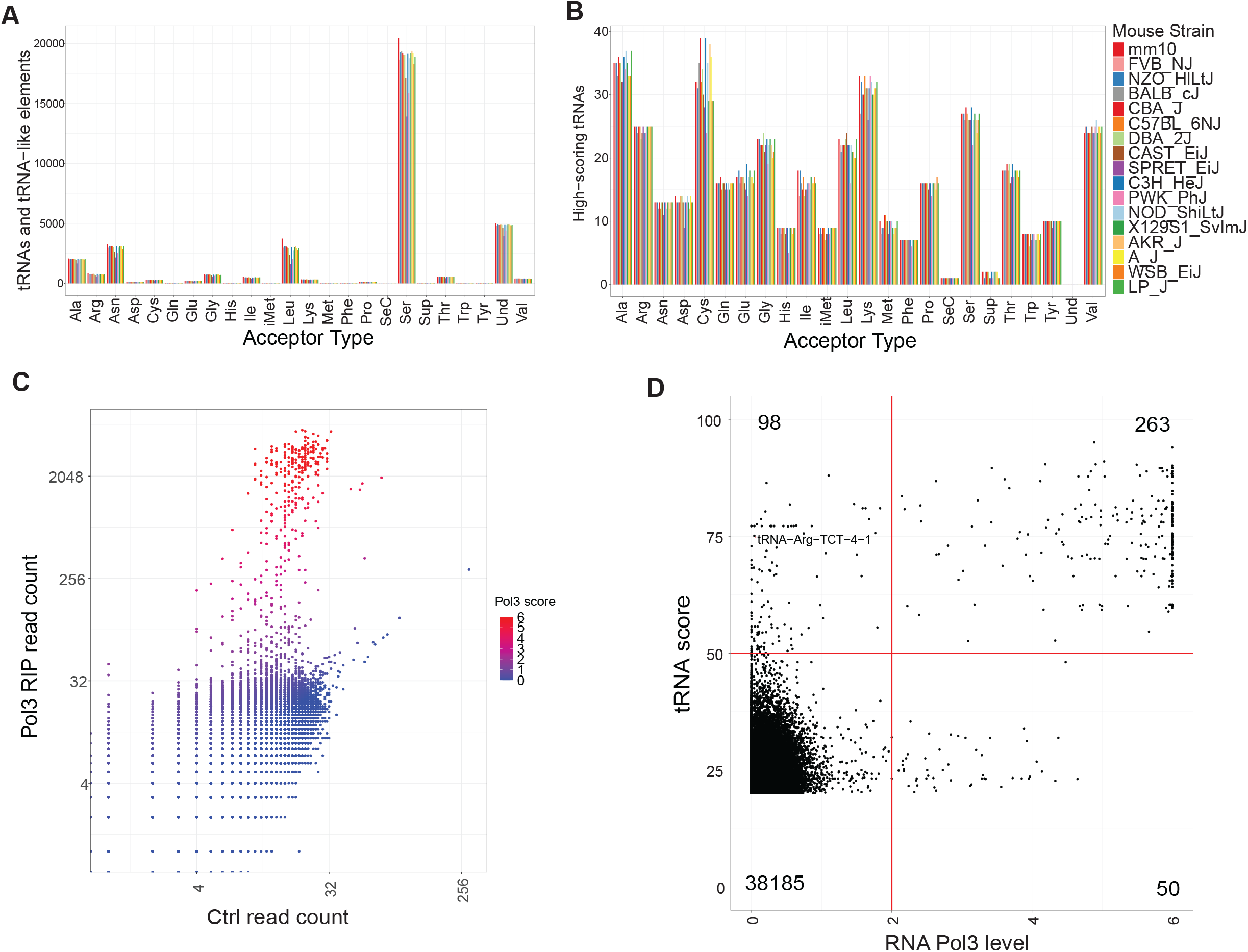
A) count of tRNA and tRNA-like elements for each assembled mouse strain and rat. B) Counts of high-scoring tRNA in each acceptor type for each strain. C) Comparison of liver RNA Pol III ChIP-seq from previously published work, comparing read count of pulldown of Pol III component RPC4 to control Igg for all tRNA-like elements, at 48 hours after liver damage. Reads are colored by a “Pol3 score” which is calculated by dividing RPC4 read count by control read count. D) Comparison of tRNA score on the y-axis (generated from tRNAscan-SE) and the Pol3 score generated from RPC4 ChIP-seq experiment x-axis. Red lines are thresholds used in subsequence analysis. E) Confusion matrix showing comparison of tRNA score with Pol3 score for each mouse tRNA.

To differentiate between inactive tRNA-like elements and active tRNA genes, we used their covariance score as calculated by tRNAscan-SE ^11^, which measures how close their sequence and structure are to known active tRNAs. To create a putative set of transcribed and functional tRNAs for the reference genome that can be easily extended to other strains, we used a tRNA covariance score cutoff of 50, which resulted in a greatly reduced set of putative transcribed tRNAs in these genomes, reducing from 39052 to 390 tRNAs (**Figure 1B**). For instance, without a tRNA score cut-off, there are 20480 Serine tRNAs and tRNA-like element genes, and this number reduces to 27 when only considering the high-scoring, potentially active tRNA genes. While there are more sophisticated approaches that attempt to detect SINE elements directly [gtRNAdb], this tRNA covariance score method is easily extendable to other strains, even those with incomplete or suboptimal assemblies. To test if our threshold of tRNA score of 50 accurately predicts actively transcribed tRNA genes, we used previously published RNA Pol III ChIP-seq data ^24^ that measured binding of the RNA Pol III subunit RPC4 to the reference mouse genome during liver regeneration after partial hepatectomy. This system allows the capture of the maximum number of tRNA genes that can be expressed under differentiated and proliferative conditions. RNA Pol III was specifically enriched in a specific set of tRNA genes, as measured by tRNA read counts from the RPC4-bound genomic regions compared to the no-antibody control (**Figure 1C**). By dividing RPC4-pulldown enriched reads by control reads, we can calculate a “Pol III enrichment score” for each gene, indicating the level of RNA Pol III binding at that locus. Comparing this RNA Pol III enrichment score to the tRNA covariance score revealed that covariance scores correlate well with RNA Pol III activity (**Figure 1D**), demonstrating that covariance scores can predict active tRNA genes. However, using an RNA Pol III enrichment score cutoff of 2 revealed 98 “false positives” genes that have a high tRNA score, but RNA Pol III does not bind to those genes. We also detected 50 “false negatives” that were low-scoring but transcribed in regenerating liver based on RNA Pol III enrichment scores. As the RNA Pol III-ChIP data is from the liver, only tRNAs active in this tissue can be identified, and a subset of “false positive” high-scoring tRNA genes may be transcribed in other tissues. For example, the brain-transcribed tRNA-Arg-TCT-4 ^17^ did not show RNA Pol III enrichment in the liver at this threshold (**Figure 1D**). Nonetheless, as most tRNA genes in the reference genome with a covariance score of 50 or more showed RNA Pol III enrichment, our analysis revealed that the tRNA score is a viable approach to predict active tRNA genes across mouse strains.

### Single-copy conserved tRNAs across mouse strains, implicating specialized function

tRNA gene sequences can occur as a single copy or as multiple identical copies spread out in the genome. To examine if tRNA copy number varied across mouse strains, we identified single-copy and multiple-copy tRNAs across the 17 mouse strains. We found that roughly half of all high-scoring mouse tRNA loci have a tRNA sequence that exists in multiple identical copies in the current reference assembly, with the other half existing as unique single-copy tRNAs (**Figure 2A**). Moreover, consistent with tRNAs being slow-evolving, these multicopy tRNAs also exist in multiple copies in the human genome (**Figure 2B)**. As expected, a comparison of different mouse strains to the reference genome revealed that tRNA copy numbers remained mostly constant between mouse strains. For instance, the dominant multicopy tRNA genes like tRNA-Cys-CGA-4 and tRNA-Gly-GCC_2 exist in identical copy number in the reference mm10 genome and FVB/NJ strain genome (**Figure 2B**). However, we also identified some tRNAs that vary in copy number across strains, for example, tRNA-Asp-GTC-1, which is present in 11 copies in mm10 and 9 copies in FVB/NJ. Across all strain comparisons, we found extreme cases where tRNA copy number can differ by up to 5 copies between any two mouse strains (**Figure 2C**). Due to the suboptimal genome assemblies of some of these strains, it is possible that some tRNA copies existing in unassembled regions will not be assessed. However, we did not observe highly variable tRNA copy numbers or changes in the dominant tRNA decoders across strains. Thus, our analysis indicates that these assemblies are accurately distinguishing single-copy and multicopy tRNAs, and that copy number varies across strains for some tRNAs.

**Figure 2.**
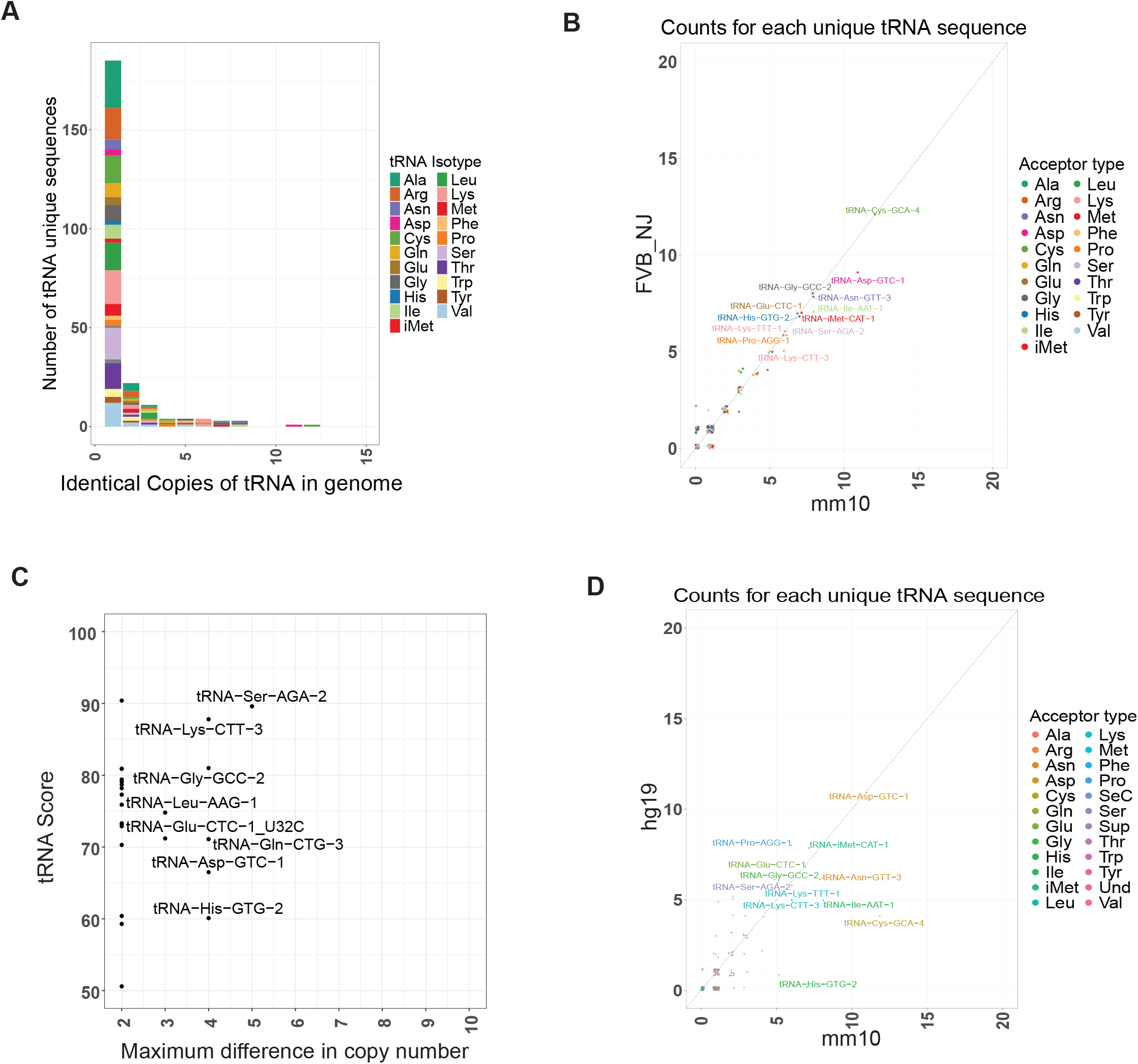
A) Histogram of tRNA copy number for all high-scoring tRNAs in the mm10 assembly. B) Comparison of tRNA copy number for high-scoring tRNAs in the mm10 assembly compared to high-scoring tRNAs in the FVB strain assembly. C) Comparison of tRNA score on the y axis compared to the difference between the maximum tRNA count and the minimum tRNA count among all strain assemblies, excluding rat.

Although multiple-copy tRNAs are generally predicted to be more likely to be transcribed compared to tRNAs that exist in a single copy ^14^, we next investigated whether single-copy tRNAs are transcribed and could therefore be involved in specialized functions. One previously discovered specialized single-copy tRNA is the brain-specific tRNA-Arg-TCT-4, discovered in the human genome ^17^. A slight variant of this tRNA exists in the mouse reference genome, also annotated as tRNA-Arg-TCT-4, differing from the human tRNA sequence by a single base (**Figure S1**). We find that the exact sequence of this human brain-specific tRNA is present in the genomes of all mouse strains, except for the reference strain (C57BL/6J), suggesting a recent change in this tRNA in the C57BL/6J strain. As in the human genome, this tRNA exists as a single copy in all mouse strain assemblies. These studies revealed that this brain-specific tRNA is tightly conserved and does not duplicate, suggesting that special-purpose tRNAs must retain their sequence to retain their specialized function and avoid duplication that may cause misregulation or excess transcript production. Single-copy tRNAs can exist either as a recent tRNA variant mutated from a multicopy version or as an ancestral tRNA variant that is conserved, possibly for some specialized purpose. To identify additional tRNAs that could be specialized single-copy tRNAs, we searched mouse genomes for other high-scoring tRNAs that existed in a single copy in all or most mouse strains. Despite the close relation of these stains, we find that most of these single-copy tRNAs are transient, existing as a single copy only in a single mouse strain (**Figure 3A**). These transient tRNAs can be either mutating or absent from other mouse strain assemblies. Notably, we identified 92 tRNAs that exist as single-copy in at least 14 mouse genomes, with their conservation making them candidates for specialized tRNAs (**Figure 3A**). Using RNA Pol III ChIP-seq data, we find that many of these single-copy tRNAs that are conserved across all mouse strains are expressed, while single-copy tRNAs that exist in fewer genomes have lower RNA Pol III enrichment (**Figure 3B**). This data implies that these single-copy tRNAs are not only conserved but are also functional, expressed tRNA genes, at least in the liver tissue. The brain-specific tRNA, here conserved as a single unique gene in all but a single assembly, was found not to be representative of this set, which is expected as it is not transcribed in the liver. To assess the conservation of single-copy tRNA genes, we examined the tRNA complement of select mammalian genomes. We examined the copy number of the 92 mouse single-copy tRNAs in the rat, human, dog, and possum genomes. Of these, 42 (45.7%) remain single-copy in rat, while 31 (33.7%) are conserved as single-copy in human. Most of the remaining tRNAs are either absent from these genomes or present in multiple copies in other species (**Figure 3C**). The non-conserved single-copy tRNA genes in the mouse may be specialized in a species-specific manner. In the set of single-copy conserved tRNAs, 9 tRNAs are conserved in a single copy in at least 15 mouse strains, as well as rat, human, dog, and possum (**Table 1**), and all except for tRNA-Arg-TCT-4 are expressed in mouse liver according to RNA Pol III ChIP-seq.

**Table 1:** Conserved single-copy tRNAs. Copy numbers of all tRNAs with a single copy in more than 15 mouse strain genomes and a single copy in the rn6 rat genome, the canFam3 dog genome, the hg19 human genome, and the monDom5 opossum genome.

| tRNA name | score | mm10 | FVB_NJ | rn6 | canFam3 | hg19 | monDom5 |
| --- | --- | --- | --- | --- | --- | --- | --- |
| tRNA-Arg-CCT-4 | 65.6 | 1 | 1 | 1 | 1 | 1 | 1 |
| tRNA-Arg-TCT-2 | 70.8 | 1 | 1 | 1 | 1 | 1 | 1 |
| tRNA-Arg-TCT-4_C50U | 78.5 | 0 | 1 | 1 | 1 | 1 | 1 |
| tRNA-Cys-GCA-2 | 81.8 | 1 | 1 | 1 | 1 | 1 | 1 |
| tRNA-Gln-TTG-1 | 72.6 | 1 | 1 | 1 | 1 | 1 | 1 |
| tRNA-Ser-CGA-2 | 89.6 | 1 | 1 | 1 | 1 | 1 | 1 |
| tRNA-Ser-TGA-1 | 94 | 1 | 1 | 1 | 1 | 1 | 1 |
| tRNA-Thr-CGT-1 | 79.6 | 1 | 1 | 1 | 1 | 1 | 1 |
| tRNA-Trp-CCA-2 | 74.3 | 1 | 1 | 1 | 1 | 1 | 1 |

**Figure 3.**
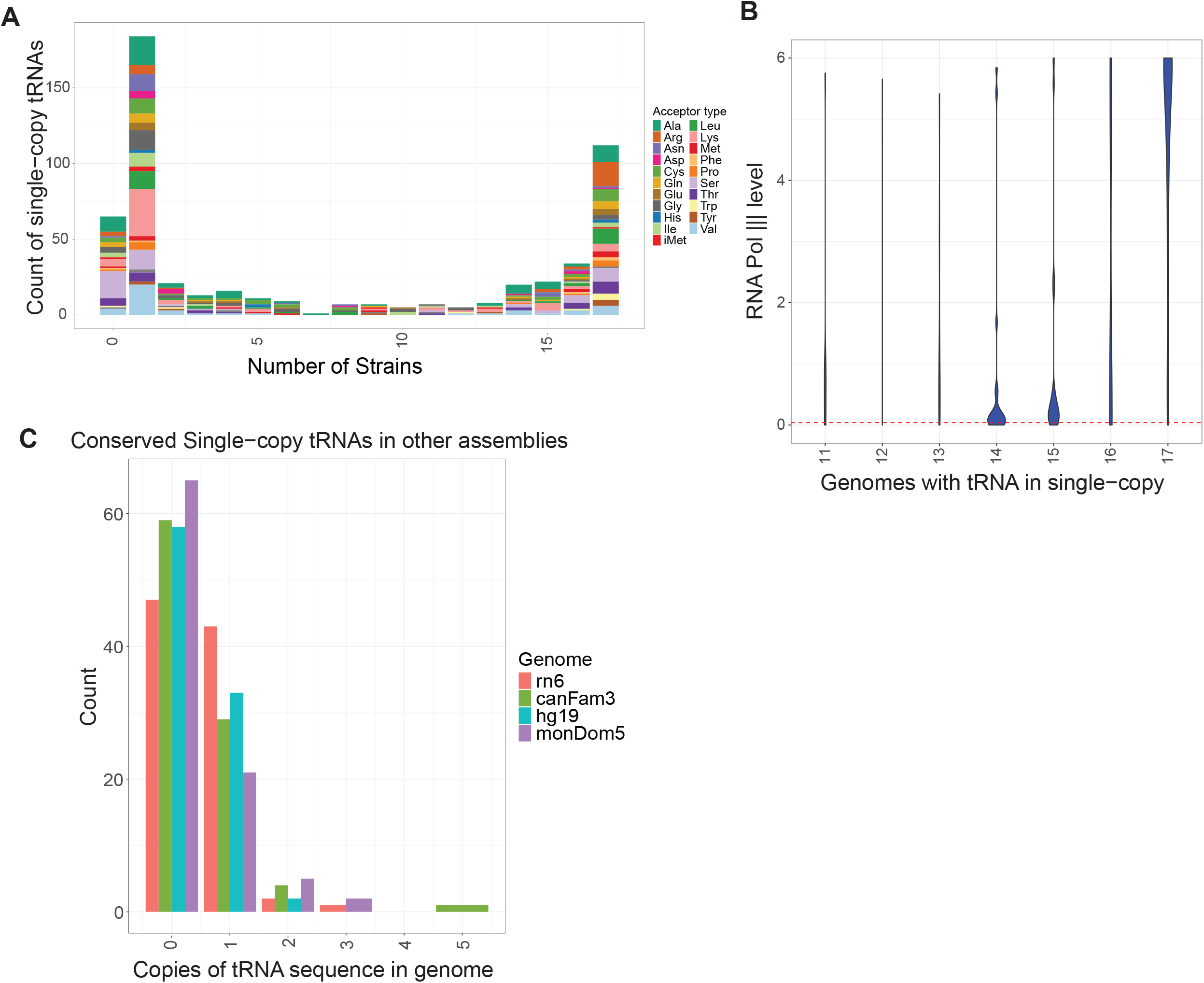
A) Histogram of the number of strains in which each unique tRNA transcript is single-copy. B) Comparison of the number of strains in which each tRNA is single-copy to the tRNA score, including only those tRNAs that exist in the mm10 assembly. These genes are colored by the Pol3 score, showing red for those genes that have a Pol 3 signal. Horizontal black line is the marker for the Pol 3 score of the brain-specific mouse tRNA, which is otherwise absent. C) Histogram of copies of each tRNA present in a single copy in at least 10 mouse strains for rat, dog, human, and possum genomes.

### Orthologous tRNAs change even in closely related mouse strains

As tRNAs exist in multiple copies in the genome, determining orthologs by sequence similarity is not possible, as it is for protein-coding genes. Therefore, we used previously published whole-genome alignment cactus graphs ^25^ to find sets of orthologous tRNAs. These cactus graph alignments include the genomes of 17 assembled mouse strains and the rat genome assembly as an outgroup, and utilizing genome sequence alignment can be used to find the orthologous region of the genome for much of these assemblies. By collecting genome alignments from annotated tRNAs or tRNA-like elements from all 17 mouse assemblies, we collected a set of orthologous tRNAs and tRNA-like elements in all mouse strains and rat (**Figure 4A**). For each tRNA gene, these ortholog sets identify its corresponding ortholog in each mouse strain and in rat, where such an ortholog can be found. Orthologs could not be assigned for all tRNAs across all strains, either because the gene was absent from the alignment or because the sequence contained ambiguities. With this method, we were able to find orthologous tRNAs or tRNA-like elements for more than 88% of all high-scoring tRNAs in all strains, with the rest missing either due to changes in the genome or incomplete assemblies. We found that high-scoring tRNA genes change in sequence even between closely related strains, with 5 tRNAs changing in sequence in the most closely related strain to the reference genome, C57BL/6NJ, and up to 32% of tRNAs changing in the most distantly related strain, Mus spretus (**Figure 4B**). The rat genome showed the most change in orthologous tRNAs compared to the reference mouse genome with only 178 (45%) of tRNAs remaining identical and 58 (15%) of tRNAs changing by at least one nucleotide.

**Figure 4.**
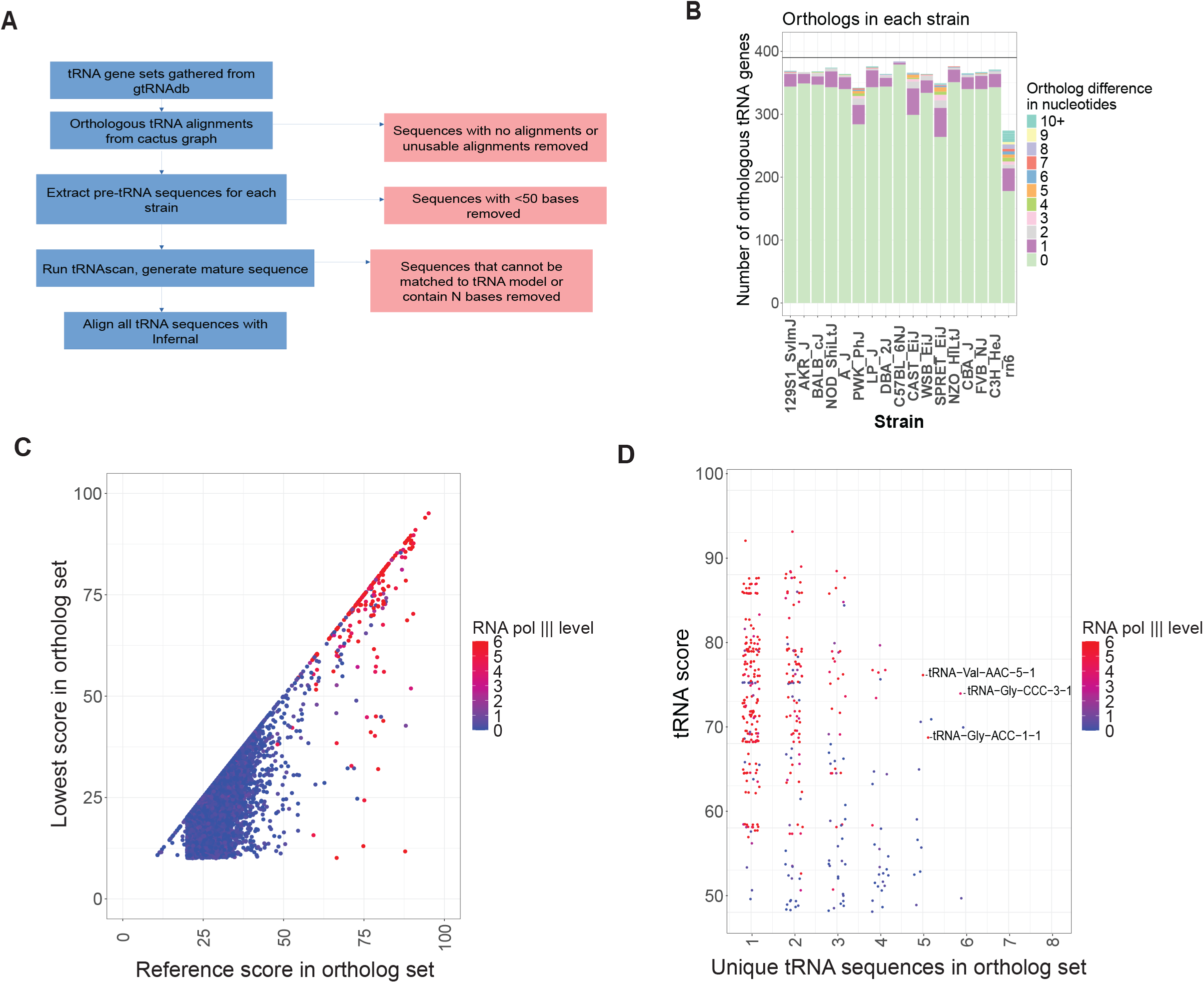
A) Flowchart of tRNA ortholog creation from mouse strain cactus graphs. Red squares show places where tRNAs are lost from the total set, either due to not having orthologs or because the ortholog cannot be found or exactly sequenced. B) Count of tRNA orthologs for high-scoring tRNA sequences in the mouse reference in each non-reference mouse strain and rat. Counts are colored by the number of mismatches present in each strain C) Comparison of reference tRNA score to the lowest score among all orthologs of that tRNA for all tRNA-like elements. Dots are colored by their Pol3 score to show the level of Pol III expression. D) Comparison of tRNA score to show the quality of tRNA to the number of unique sequences to show variation across all mouse strains. Dots are colored by their Pol3 score to show the level of RNA Pol III expression. tRNAs expressed in reference with 5 or more sequences are labeled.

These differences between related genomes include tRNAs for which an orthologous sequence can be found but appear to be nonviable tRNAs as measured by their tRNA scores, which measure the loss of structure or ability to be transcribed. Comparing tRNAs in other strains to the reference, we find 14 tRNAs that are both high-scoring and confirmed as transcribed in the reference mouse genome by RNA Pol III ChIP-seq that appear to have lost their function in other mouse strains, due to sequence changes in specific regions of the tRNA (promoters or other key structural features) (**Figure 4C**). While some tRNAs remain identical in sequence across all mouse strains, some tRNAs change more than once. For instance, we found that some tRNA ortholog sets include up to 6 unique variants among their orthologs (**Figure 4D**). Importantly, this also includes tRNAs for which RNA Pol III ChIP-seq transcription is confirmed, such as tRNA-Val-AAC-5-1, tRNA-Gly-CCC-3-1, and tRNA-Gly-ACC-1-1 (**Figure 4D** and **Figure S2A-C**).

### Gain and loss of tRNA genes in related mouse genomes

We can also identify tRNA genes that have been recently added or are missing in the genomes of different mouse strains in comparison to the mouse reference genome, to study the evolution of tRNA genes. To do this, we counted the number of high-scoring tRNAs in each ortholog set, with putatively transcribed tRNAs in almost all strains being likely deletions in the remaining strains, and those present in very few strains being new insertions. We found the pattern of strain counts consistent with this, with ortholog set counts tending towards one strain or all strains, with few ortholog sets in the moderate range of 5-10 strains (**Figure 5A**). Using this approach, we can look at the set of tRNAs that exist in the reference genome but are not conserved (**Figure 5B**). Our analysis revealed that this set includes many tRNAs that, while high scoring, have low RNA Pol III score, indicating they are either not transcribed or transcribed in only some conditions.

**Figure 5.**
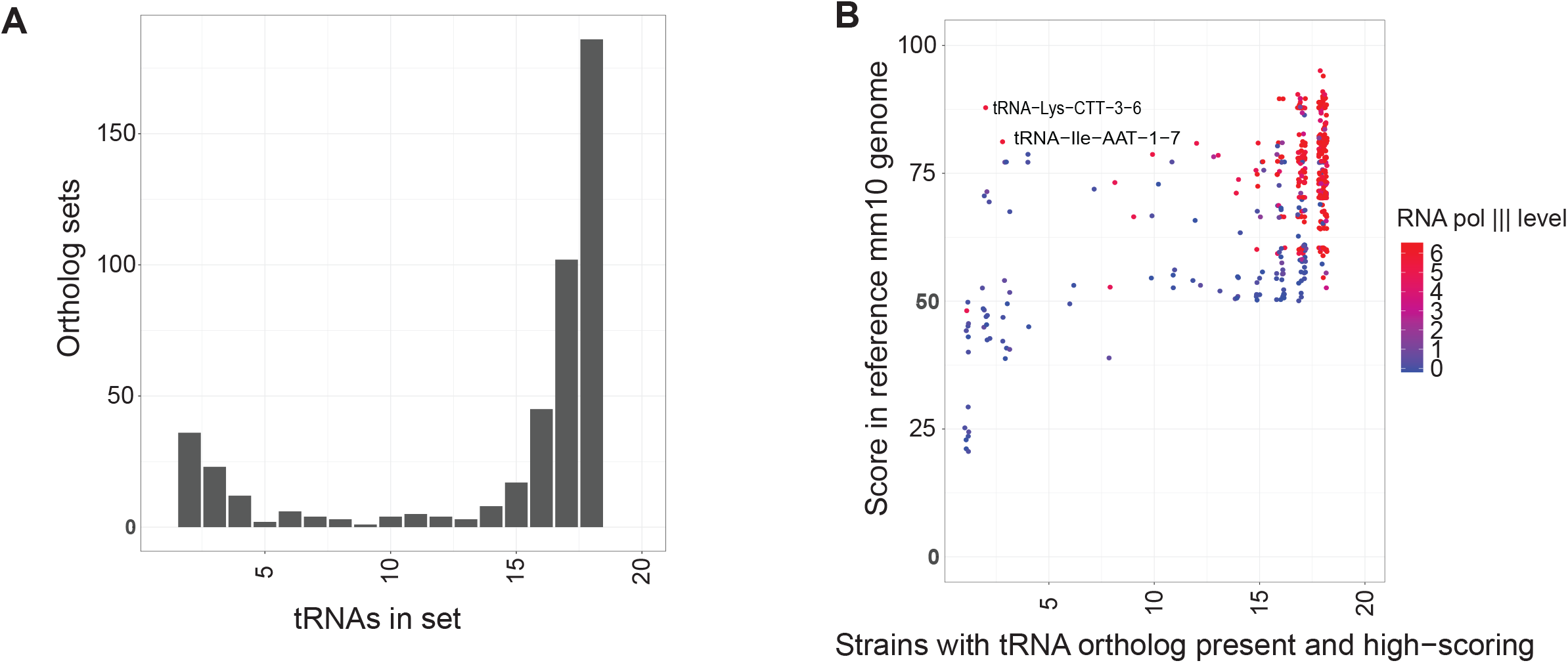
A) Histogram of the number of mouse strains included in each predicted tRNA ortholog set. Only high-scoring (>50) tRNAs are counted. B) Scatter plot comparing tRNA score of reference mm10 tRNAs with the number of high-scoring orthologs for each ortholog set. Dots are colored by their Pol3 score to show the level of RNA Pol III expression.

## Discussion

Our study provides a systematic analysis of tRNA gene conservation, copy number variation, and transcriptional activity across 17 mouse strains, using a combination of covariance scoring and RNA polymerase III ChIP-seq validation. While tRNA genes are among the most conserved features of mammalian genomes ^4-6^, our results demonstrate that significant variability exists in both sequence and function, even among closely related strains. This indicates that the tRNA complement is not as static as previously assumed, but rather subject to dynamic evolutionary changes that may have functional consequences for translation and gene regulation.

By integrating sequence-based covariance scores with Pol III ChIP-seq data, we demonstrate that covariance scoring provides a useful and reasonably accurate predictor of transcriptionally active tRNAs. Although some false positives and negatives remain, some of these discrepancies can be attributed to tissue specificity, as our Pol III-Chip-seq data were derived from liver tissues. For example, the brain-specific tRNA-Arg-TCT-4 was absent from liver datasets but remains highly conserved and expressed in neuronal contexts. These findings highlight the importance of integrating structural predictions with tissue- and condition-specific transcriptional data to fully capture the functional repertoire of tRNA genes.

We also find that tRNA copy numbers are largely stable across mouse strains, consistent with the strong evolutionary constraints on tRNA dosage and decoding balance. Nevertheless, some tRNAs exhibit copy number differences of up to five copies between strains. Although tRNAs are often considered redundant elements in translational regulation, prior work has demonstrated that certain single-copy, uniquely occurring tRNAs can carry specialized functions ^17^. In our analysis across multiple mouse strains, we found that while many single-copy tRNAs are not conserved, a subset remains conserved and thus represent strong candidates for specialized roles. Interestingly, these conserved single-copy tRNAs generally do not follow the pattern of the well-characterized brain-specific tRNA, but instead show RNA Pol III ChIP-seq signatures of expression in liver. The evolutionary pressure to maintain these tRNAs as single, unique copies is not yet clear, but their conservation suggests functional importance. That all but one are expressed in liver does suggest that tissue specificity is not the primary form of specialization. Possible roles include contributing to the translation of a restricted set of genes, functioning in a conditionally regulated manner where they are expressed or silenced depending on context, or serving as a source of regulatory tRNA fragments with distinct biological functions.

Finally, our ortholog analyses revealed that actively transcribed tRNAs can be gained, lost, or altered in sequence even among closely related strains. Some transcribed orthologous sets showed multiple sequence variants, while others lost predicted structural integrity despite being transcribed in the reference genome. These changes underscore that while the tRNA pool is highly conserved overall, subsets of tRNAs are evolving rapidly, with potential implications for translational efficiency, codon bias adaptation, and phenotypic diversity across strains. Together, our results provide a resource for understanding the evolution of the tRNA repertoire in mammals and highlight the need to consider tRNA variation in studies of gene regulation, genome evolution, and strain-specific phenotypes.

## Methods

### Generating mouse ortholog sets

Mouse tRNA annotations for reference tRNA and all sequenced strains were gathered through the genomic tRNA database ^22^. These annotations were compared to a mouse strain cactus graph alignment ^25^. Multi-strain sequence alignments of all tRNAs and tRNA-like elements with 20 flanking bases were calculated for all tRNAs in all other genomes using the liftover ^26^ tool, removing duplicates and merging regions to remove discontinuities. The set of unique sequences generated with this was run through tRNAscan-SE ^11^ with the minimum score set to 10. The output of that was used to generate the mature tRNA sequence for each of these by removing the intron, adding the CCA tail, and adding the post-transcriptional “G” base to histidine tRNAs. Sequences with N bases were also discarded at this point to remove ambiguity. tRNAscan-SE scores were also taken at this point to evaluate the quality of the tRNA, and sequences with no tRNAscan-SE result were discarded. These sequences were then aligned using infernal cmalign ^10^ to a mature eukaryotic tRNA sequence covariance model ^11^. Sequence differences were calculated for each tRNA to all other high-scoring tRNAs it is orthologous to in addition to the “quality set” of high-scoring mouse reference genome tRNAs. tRNAs not present in the set were classified as “missing” if no tRNA alignment could be found in that strain and “pseudo” if no tRNA-like sequence could be detected by tRNAscan-SE.

### Mouse RNA polymerase III chip-seq analysis

Mouse RNA Pol III chip-seq data were generated in a previous study that examined polymerase data in regenerating and control mouse liver pulled down with the RPC4 subunit ^24^. Reads from this were mapped to the mouse mm10 reference genome with bowtie2 ^27^. Reads for each sample were counted within 50 bases of a specific tRNA to generate read counts for that tRNA. The Pol3 score was calculated by dividing the number of reads in the regenerating liver pulldown by the control liver, setting all scores < 0 to zero and all scores > 6 to 6.

## Supplementary Material

**Supplementary Figure S1:** Sequence alignment of the tRNA-Arg-TCT-4 tRNA between mm10 mouse genome, hg19 human genome, and FVB mouse genome. Base position that differs is highlighted in yellow.

**Supplementary Figure S2:** Sequence alignment of the ortholog sets for tRNA-Gly-ACC-1-1, tRNA-Gly-CCC-3-1, and tRNA-Val-AAC-5-1 tRNA genes. Showing the sequence for the orthologous gene in each strain. Sequences are color-coded such that identical tRNAs share the same color, allowing differences between strains to be readily visualized.

**Supplementary Table S1:** Table of tRNA sequences for each tRNA or tRNA-like sequence found in any mouse strain. Amino acid, anticodon, and scores are predicted from tRNAscan-SE. Names correspond to the mm10 reference names where possible, and new names are created for sequences not present in the mm10 reference tRNA set. Number of copies of a given tRNA shown for each mouse strain and for the rn6 rat genome.

**Supplementary Table S2:** Table of ortholog sets for tRNAs and tRNA-like elements. ‘NA’ indicates that no orthologous tRNA-like element was identified. tRNA names correspond to the sequences listed in Table S1. When applicable, the ortholog set name includes the locus identifier of the gene in the reference mm10 strain; if not, the locus identifier from another strain is provided.

## Notes

### Competing Interest Statement

The authors have declared no competing interest.

### Funding

- National Institutes of Health: 1DP2AG066622-01

## References

1. Holmes, A.D., Chan, P.P., Chen, Q., Ivanov, P., Drouard, L., Polacek, N., Kay, M.A., and Lowe, T.M. (2023). A standardized ontology for naming tRNA-derived RNAs based on molecular origin. Nat Methods 20, 627–628. 10.1038/s41592-023-01813-2.

2. Anderson, P., and Ivanov, P. (2014). tRNA fragments in human health and disease. FEBS letters 588, 4297–4304. 10.1016/j.febslet.2014.09.001.

3. Polacek, N., and Ivanov, P. (2020). The regulatory world of tRNA fragments beyond canonical tRNA biology. Rna Biol 17, 1057–1059. 10.1080/15476286.2020.1785196.

4. Tang, D.T., Glazov, E.A., McWilliam, S.M., Barris, W.C., and Dalrymple, B.P. (2009). Analysis of the complement and molecular evolution of tRNA genes in cow. BMC Genomics 10, 188. 10.1186/1471-2164-10-188.

5. Westhof, E., Thornlow, B., Chan, P.P., and Lowe, T.M. (2022). Eukaryotic tRNA sequences present conserved and amino acid-specific structural signatures. Nucleic acids research 50, 4100–4112. 10.1093/nar/gkac222.

6. Kutter, C., Brown, G.D., Goncalves, A., Wilson, M.D., Watt, S., Brazma, A., White, R.J., and Odom, D.T. (2011). Pol III binding in six mammals shows conservation among amino acid isotypes despite divergence among tRNA genes. Nat Genet 43, 948–955. 10.1038/ng.906.

7. Fujishima, K., and Kanai, A. (2014). tRNA gene diversity in the three domains of life. Front Genet 5, 142. 10.3389/fgene.2014.00142.

8. Zhang, J. (2024). Recognition of the tRNA structure: Everything everywhere but not all at once. Cell Chem Biol 31, 36–52. 10.1016/j.chembiol.2023.12.008.

9. Sprinzl, M., Grueter, F., Spelzhaus, A., and Gauss, D.H. (1980). Compilation of tRNA sequences. Nucleic acids research 8, r1–r22.

10. Nawrocki, E.P., and Eddy, S.R. (2013). Infernal 1.1: 100-fold faster RNA homology searches. Bioinformatics 29, 2933–2935. 10.1093/bioinformatics/btt509.

11. Chan, P.P., Lin, B.Y., Mak, A.J., and Lowe, T.M. (2021). tRNAscan-SE 2.0: improved detection and functional classification of transfer RNA genes. Nucleic acids research 49, 9077–9096. 10.1093/nar/gkab688.

12. Sharp, S., DeFranco, D., Dingermann, T., Farrell, P., and Soll, D. (1981). Internal control regions for transcription of eukaryotic tRNA genes. Proc Natl Acad Sci U S A 78, 6657–6661. 10.1073/pnas.78.11.6657.

13. Canella, D., Bernasconi, D., Gilardi, F., LeMartelot, G., Migliavacca, E., Praz, V., Cousin, P., Delorenzi, M., Hernandez, N., and Cycli, X.C. (2012). A multiplicity of factors contributes to selective RNA polymerase III occupancy of a subset of RNA polymerase III genes in mouse liver. Genome Res 22, 666–680. 10.1101/gr.130286.111.

14. Thornlow, B.P., Armstrong, J., Holmes, A.D., Howard, J.M., Corbett-Detig, R.B., and Lowe, T.M. (2020). Predicting transfer RNA gene activity from sequence and genome context. Genome Res 30, 85–94. 10.1101/gr.256164.119.

15. Graczyk, D., Ciesla, M., and Boguta, M. (2018). Regulation of tRNA synthesis by the general transcription factors of RNA polymerase III - TFIIIB and TFIIIC, and by the MAF1 protein. Biochim Biophys Acta Gene Regul Mech 1861, 320–329. 10.1016/j.bbagrm.2018.01.011.

16. Kapur, M., Molumby, M.J., Guzman, C., Heinz, S., and Ackerman, S.L. (2024). Cell-type-specific expression of tRNAs in the brain regulates cellular homeostasis. Neuron 112, 1397–1415 e1396. 10.1016/j.neuron.2024.01.028.

17. Ishimura, R., Nagy, G., Dotu, I., Zhou, H., Yang, X.L., Schimmel, P., Senju, S., Nishimura, Y., Chuang, J.H., and Ackerman, S.L. (2014). RNA function. Ribosome stalling induced by mutation of a CNS-specific tRNA causes neurodegeneration. Science 345, 455–459. 10.1126/science.1249749.

18. Goodenbour, J.M., and Pan, T. (2006). Diversity of tRNA genes in eukaryotes. Nucleic acids research 34, 6137–6146. 10.1093/nar/gkl725.

19. Iben, J.R., and Maraia, R.J. (2014). tRNA gene copy number variation in humans. Gene 536, 376–384. 10.1016/j.gene.2013.11.049.

20. Naidoo, K., Steenkamp, E.T., Coetzee, M.P., Wingfield, M.J., and Wingfield, B.D. (2013). Concerted evolution in the ribosomal RNA cistron. PloS one 8, e59355. 10.1371/journal.pone.0059355.

21. Amstutz, H., Munz, P., Heyer, W.D., Leupoid, U., and Kohli, J. (1985). Concerted evolution of tRNA genes: intergenic conversion among three unlinked serine tRNA genes in S. pombe. Cell 40, 879–886. 10.1016/0092-8674(85)90347-2.

22. Chan, P.P., and Lowe, T.M. (2016). GtRNAdb 2.0: an expanded database of transfer RNA genes identified in complete and draft genomes. Nucleic acids research 44, D184–189. 10.1093/nar/gkv1309.

23. Veniaminova, N.A., Vassetzky, N.S., and Kramerov, D.A. (2007). B1 SINEs in different rodent families. Genomics 89, 678–686. 10.1016/j.ygeno.2007.02.007.

24. Yeganeh, M., Praz, V., Carmeli, C., Villeneuve, D., Rib, L., Guex, N., Herr, W., Delorenzi, M., Hernandez, N., and Cycli, X.c. (2019). Differential regulation of RNA polymerase III genes during liver regeneration. Nucleic acids research 47, 1786–1796. 10.1093/nar/gky1282.

25. Paten, B., Diekhans, M., Earl, D., John, J.S., Ma, J., Suh, B., and Haussler, D. (2011). Cactus graphs for genome comparisons. J Comput Biol 18, 469–481. 10.1089/cmb.2010.0252.

26. Perez, G., Barber, G.P., Benet-Pages, A., Casper, J., Clawson, H., Diekhans, M., Fischer, C., Gonzalez, J.N., Hinrichs, A.S., Lee, C.M., et al. (2025). The UCSC Genome Browser database: 2025 update. Nucleic acids research 53, D1243–D1249. 10.1093/nar/gkae974.

27. Langmead, B., and Salzberg, S.L. (2012). Fast gapped-read alignment with Bowtie 2. Nat Methods 9, 357–359. 10.1038/nmeth.1923.

