## Supplementary figures for "Tracking tRNA evolution in closely related mouse strains"

#### Supplementary Figure 1

```
mm10    tRNA-Arg-TCT-4-1     GTCTCTGTGGCGCAATGGACGAGCGCGCTGGACTTCTAATCCAGAGGTTCTGGGTTCGAGTCCCGGCAGAGATGCCA
hg19    tRNA-Arg-TCT-4-1     GTCTCTGTGGCGCAATGGACGAGCGCGCTGGACTTCTAATCCAGAGGTTCCGGGTTCGAGTCCCGGCAGAGATGCCA
FVB_NJ  tRNA-Arg-TCT-4-1     GTCTCTGTGGCGCAATGGACGAGCGCGCTGGACTTCTAATCCAGAGGTTCCGGGTTCGAGTCCCGGCAGAGATGCCA
```

### Supplementary Figure 2

**A** tRNA-Gly-ACC-1-1

```
mm10          GTTTCCGTAGTGTAGTGGTTAGCGCGTTCGCCTACCAAAGCGAAAGGTCCCCGGTTCGAAACCGGGCGGAAACACCA
FVB_NJ        GTTTCCGTAGTGTAGTGGTTAGCGCGTTCGCCTACCAAAGCGAAAGGTCCCCGGTTCGAAACCGGGCGGAAACACCA
NZO_HlLtJ     GTTTCCGTAGTGTAGTGGTTAGCACGTTCGCCTACCAAAGCGAAAGGTCCCCGGTTCGAAACCGGGCGGAAACACCA
BALB_cJ       GTTTCCGTAGTGTAGTGGTTAGCGCGTTCGCCTACCAAAGCGAAAGGTCCCCGGTTCGAAACCGGGCGGAAACACCA
CBA_J         GTTTCCGTAGTGTAGTGGTTAGCGCGTTCGCCTACCAAAGCGAAAGGTCCCCGGTTCGAAACCGGGCGGAAACACCA
C57BL_6NJ     GTTTCCGTAGTGTAGTGGTTAGCGCGTTCGCCTACCAAAGCGAAAGGTCCCCGGTTCGAAACCGGGCGGAAACACCA
DBA_2J        GTTTCCGTAGTGTAGTGGTTAGCGCGTTCGCCTACCAAAGCGAAAGGTCCCCGGTTCGAAACCGGGCGGAAACACCA
CAST_EiJ      GTTTCCGTAGTGTAGTGGTTAGCGCGTTCGCCTACCAAAGCGAAAGGTCCCCGGTTCGAAACCGGGCGGAAACACCA
SPRET_EiJ     GTTTCCGTAGTGTAGTGGTTATCACGTTCGCCTA-ACACGCGAAAGGTCCCCGGTTCAAAACCGGGCGGAAACACCA
C3H_HeJ       GTTTCCGTAGTGTAGTGGTTAGCGCGTTCGCCTACCAAAGCGAAAGGTCCCCGGTTCGAAACCGGGCGGAAACACCA
PWK_PhJ       GTTTCCGTAGTGTAGTGGTTATCACGTTCGCCTA-ACACGCGAAAGGTCCCCGGTTCGAAACCGGGCGGAAACACCA
NOD_ShiLtJ    GTTTCCGTAGTGTAGTGGTTAGCGCGTTCGCCTACCAAAGCGAAAGGTCCCCGGTTCGAAACCGGGCGGAAACACCA
X129S1_SvImJ  GTTTCCGTAGTGTAGTGGTTAGCGCGTTCGCCTACCAAAGCGAAAGGTCCCCGGTTCGAAACCGGGCGGAAACACCA
AKR_J         GTTTCCGTAGTGTAGTGGTTAGCACGTTCGCCTACCAAAGCGAAAGGTCCCCGGTTCGAAACCGGGCGGAAACACCA
A_J           GTTTCCGTAGTGTAGTGGTTAGCGCGTTCGCCTACCAAAGCGAAAGGTCCCCGGTTCGAAACCGGGCGGAAACACCA
WSB_EiJ       GTTTCCGTAGTGTAGTGGTTAGCACGTTCGCCTACCAAAGCGAAAGGTCCCCGGTACGAAACCGGGCGGAAACACCA
LP_J          GTTTCCGTAGTGTAGTGGTTAGCGCGTTCGCCTACCAAAGCGAAAGGTCCCCGGTTCGAAACCGGGCGGAAACACCA
rn6           GTTTCCGTAGTGTAGTGGTTATCACGTTCGCCTA-ACACGCGAAAGGTCCCCGGTTCGAAACCGGGCGGAAACACCA
```

**B** tRNA-Gly-CCC-3-1

```
mm10          GCATTGGTGGTTCAATGGTAGAATTCTCGCCTCCCACGCGGGTGACCCGGGTTCGATTCCCGGCCAATGCACCA
FVB_NJ        GCATTGGTGGTTCAATGGTAGAATTCTCGCTTCCCACGCGGGTGACCCGGGTTCGATTCCCGGCCAATGCACCA
NZO_HlLtJ     GCATTGGTGGTTCAATGGTAGAATTCTCGCCTCCCATGCGGGTGACCTGGGTTCGATTCCCGGCCAATGCACCA
BALB_cJ       GCATTGGTGGTTCAATGGTAGAATTCTCGCCTCCCATGCGGGTGACCTGGGTTCGATTCCCGGCCAATGCACCA
CBA_J         GCATTGGTGGTTCAATGGTAGAATTCTCGCTTCCCACGCGGGTGACCCGGGTTCGATTCCCGGCCAATGCACCA
C57BL_6NJ     GCATTGGTGGTTCAATGGTAGAATTCTCGCCTCCCACGCGGGTGACCCGGGTTCGATTCCCGGCCAATGCACCA
DBA_2J        GCATTGGTGGTTCAATGGTAGAATTCTCGCCTCCCACGCGGGTGACCCGGGTTCGATTCCCGGCCAATGCACCA
CAST_EiJ      GCATTGGTGGTTCAATGGTAGAATTCTTGCCTCCCACGCGGGTGACCCGGGTTCGATTCCCGGCCAATGCACCA
SPRET_EiJ     GCATTGGTGGTTCAATGGTAGAATTCTCGCCTCCCACGCGGGTGACCCGGGTTCGATTCCCGGCCAATGCACCA
C3H_HeJ       GCATTGGTGGTTCAATGGTAGAATTCTCGCTTCCCACGCGGGTGACCCGGGTTCGATTCCCGGCCAATGCACCA
PWK_PhJ       GCTTTGGTGGTTCAATGGTAGAATTCTCGCCTCCCACGCGGGTGACCCGGGTTCGATTCCCGGCCAATGCACCA
NOD_ShiLtJ    GCATTGGTGGTTCAATGGTA---TTCTCGCTTCCCACGCGGGTGACCCGGGTTCGATTCCCGGCCAATGCACCA
X129S1_SvImJ  GCATTGGTGGTTCAATGGTAGAATTCTCGCCTCCCACGCGGGTGACCCGGGTTCGATTCCCGGCCAATGCACCA
AKR_J         GCATTGGTGGTTCAATGGTAGAATTCTCGCCTCCCACGCGGGTGACCCGGGTTCGATTCCCGGCCAATGCACCA
A_J           GCATTGGTGGTTCAATGGTAGAATTCTCGCTTCCCACGCGGGTGACCCGGGTTCGATTCCCGGCCAATGCACCA
WSB_EiJ       GCATTGGTGGTTCAATGGTAGAATTCTCGCCTCCCACGCGGGTGACCCGGGTTCGATTCCCGGCCAATGCACCA
LP_J          GCATTGGTGGTTCAATGGTAGAATTCTCGCCTCCCACGCGGGTGACCCGGGTTCGATTCCCGGCCAATGCACCA
rn6           GCATTGGTGGTTCAATGGTAGAATTCTCGCCTCCCACGCGGGTGACCCGGGTTCGATTCCCGGCCAATGCACCA
```

**C** tRNA-Val-AAC-5-1

```
mm10          GTTTCCGTAGTGTAGTGGTTATCACATTCGCCTAACACGCGAAAGGTCCCCGGTTCGAAACCGGGCGGAAACACCA
FVB_NJ        GTTTCCGTAGTGTAGTGGTTATCACATTCGCCTAACACGCGAAAGGTCCCCGGTTCGAAACCGGGCGGAAACACCA
NZO_HlLtJ     GTTTCCGTAGTGTAGTGGTTATCACATTCGCCTAACACGCGAAAGGTCCCCGGTTCGAAACCGGGCGGAAACACCA
BALB_cJ       GTTTCCGTAGTGTAGTGGTTATCACATTCGCCTAACACGCGAAAGGTCCCCGGTTCGAAACCGGGCGGAAACACCA
CBA_J         GTTTCCGTAGTGTAGTGGTTATCACATTCGCCTAACACGCGAAAGGTCCCCGGTTCGAAACCGGGCGGAAACACCA
C57BL_6NJ     GTTTCCGTAGTGTAGTGGTTATCACATTCGCCTAACACGCGAAAGGTCCCCGGTTCGAAACCGGGCGGAAACACCA
DBA_2J        GTTTCCGTAGTGTAGTGGTTATCACATTCGCCTAACACGCGAAAGGTCCCCGGTTCGAAACCGGGCGGAAACACCA
CAST_EiJ      GTTTCCGTAGTGTAGTGGTTATCACATTCGCCTAACACGCGAAAGGTCCCCGGTTCGAAACCGGGCGGAAACACCA
SPRET_EiJ     GTTTCCGTAGTGTAGTGGTTATCACATTCGCCTAACACGCGAAAGGTCCCCGGTTCGAAACCGGGCGGAAACACCA
C3H_HeJ       GTTTCCGTAGTGTAGTGGTTATCACATTCGCCTAACACGCGAAAGGTCCCCGGTTCGAAACCGGGCGGAAACACCA
PWK_PhJ       GTTTCCGTAGTGTAGTGGTTATCACATTCGCCTAACACGCGAAAGGTCCCCGGTTCGAAACCGGGTGGAAACACCA
NOD_ShiLtJ    GTTTCCGTAGTGTAGTGGTTATCACATTCGCCTAACACGCGAAAGGTCCCCGGTTCGAAACCGGGCGGAAACACCA
X129S1_SvImJ  GTTTCCGTAGTGTAGTGGTTATCACATTCGCCTAACACGCGAAAGGTCCCCGGTTCGAAACCGGGCGGAAACACCA
AKR_J         GTTTCCGTAGTGTAGTGGTTATCACATTCGCCTAACACGCGAAAGGTCCCCGGTTCGAAACCGGGCGGAAACACCA
A_J           GTTTCCGTAGTGTAGTGGTTATCACATTCGCCTAACACGCGAAAGGTCCCCGGTTCGAAACCGGGCGGAAACACCA
WSB_EiJ       GTTTCCGTAGTGTAGTGGTTATCACATTCGCCTAACACGCGAAAGGTCCCCGGCCCGAAACCGGGCGGAAACACCA
LP_J          GTTTCCGTAGTGTAGTGGTTATCACATTCGCCTAACACGCGAAAGGTCTCCGGTTCGAAACCGGGCGGAAACACCA
rn6           GTTTCCGTAGTGTAGTGGTTATCACATTCGCCTAACACGCGAAAGGTCCCCGGTTCGAAACCGGGCGGGAACACCA
```
